# Divergent selection and plasticity shape reproductive isolation at the onset of speciation

**DOI:** 10.1101/2024.09.23.614431

**Authors:** Benjamin J. M. Jarrett, Philip A. Downing, Erik I. Svensson

## Abstract

Reproductive isolation is a key process during speciation, but the factors that shape its evolution during the early stages of speciation remain largely unknown. Using a meta-analysis of 34 experimental speciation studies, we show that populations that experience divergent selection evolved stronger reproductive isolation compared to populations that evolved in similar environments, consistent with ecological speciation theory. However, and contrary to predictions, reproductive isolation did not increase with the number of generations. We present evidence that phenotypic plasticity could play a role in explaining these results, as divergent environments induce an increase in reproductive isolation in the first few generations. Our results highlight that adaptation to different environments in conjunction with plasticity can lead to a rapid increase in reproductive isolation at the beginning of speciation.

## Main text

The evolution of reproductive isolation is a crucial process in speciation (*1*). Rooted in the Biological Species Concept (*2, 3*), reproductive isolation can be viewed as an emergent property of the interaction between two populations that determines the likelihood of gene flow between them, though how to measure it and what it means for speciation is debated (*4–7*). Populations can be reproductively isolated from one another if they occupy different habitats (*8, 9*), if mating preferences are biased to certain genotypes or phenotypes (*10*), if hybrids are unfit (*11*), or if hybrids are not produced at all (*12, 13*). The joint action of multiple isolating barriers can quickly result in complete reproductive isolation between populations (*14, 15*). Despite being central to speciation, the mechanisms driving the evolution of reproductive isolation are still debated.

A long-standing explanation for the evolution of reproductive isolation is the ecological speciation model (*16–18*). This views that reproductive isolation evolves as a by-product of natural selection operating in different environments (*16, 17*). Adaptation in response to divergent selection imposed by different environments can result in greater genetic divergence between populations than populations experiencing the same selective pressures, and an increased likelihood that barrier loci hitchhike with the selected loci, and/or that selected loci are pleiotropic and also contribute to reproductive isolation (*17, 18*). With its roots in Darwin’s work (*19*), research during the last three decades has revealed several examples in some well-studied taxa, which strongly implicate a key role of divergent selection in speciation (*18, 20–22*).

But how general of a mechanism is ecological speciation? An alternative mechanism, also driven by selection, is mutation-order speciation, whereby populations experience similar selection pressures, and reproductive isolation evolves due to the random chance of different mutations being fixed in the different populations (*22*). A recent comparative study has suggested that mutation-order speciation is the dominant speciation mechanism in vertebrates and that ecological speciation could only explain a minority of speciation events (*23*). Since sister species were about ten times more likely to be phenotypically similar to each other, and since the phenotypes were ecologically relevant traits, the authors interpreted their results as evidence for sister species evolving in similar environments (*23*).

While comparative analyses are a powerful way to illuminate the processes that underpin the generation of new species (*5, 23–30*), they are not our only tool for studying speciation. Experimental evolution of reproductive isolation is a complementary tool, starting from the opposite end of the speciation continuum (*16, 31, 32*). Experimental evolution enables researchers to causally test mechanisms behind the evolution of reproductive isolation by dividing a single population into different replicates, imposing different experimental regimes to these replicates, and then estimating reproductive isolation after any number of generations (Figure 1A and B). While experimental evolution cannot replicate the entire speciation continuum, rapid adaptation and speciation often relies on standing genetic variation (*33, 34*), meaning that the early phase of speciation can be mimicked in the laboratory (*16*). Laboratory speciation experiments started in the 1950s, testing mechanisms of speciation, such as reinforcement, divergence with gene flow, bottlenecks, and the allopatric models of speciation. These early studies were synthesised by Rice and Hostert (*16*) in a now classic review. One conclusion from their synthesis was that divergent selection promoted the evolution of reproductive isolation via pleiotropy and/or genetic hitchhiking (*16*). This conclusion stimulated much later research into the role of divergent selection to promote reproductive isolation and provided a conceptual grounding for the theory of ecological speciation (*17, 18*). Lacking from Rice and Hostert (*16*), however, was a quantitative analysis of the strength reproductive isolation between populations evolving in divergent environments relative to that between populations evolving in similar environments. We fill this gap using a formal meta-analysis to quantitatively assess the effect of divergent selection in promoting reproductive isolation, and examine the factors that influence its evolution.

**Figure 1.**
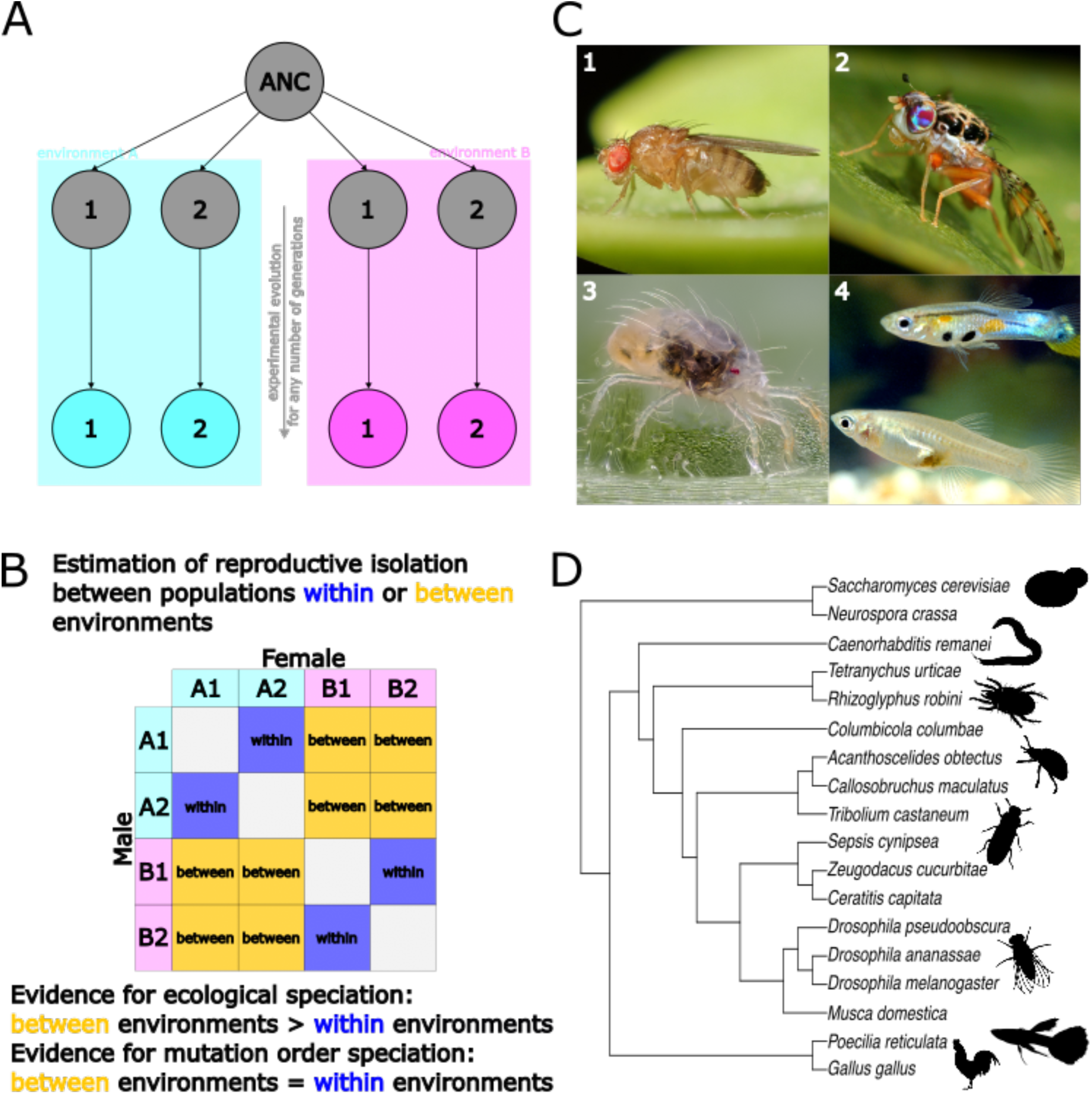
An overview of the type of experiment and effect sizes, and species included in the dataset. (**A**) An outline of the experimental design we sought: a single ancestral (ANC) population is split into two or more replicate populations that are exposed to different environments or selective regimes (represented by blue and pink) and allowed to experimentally evolve in these environments for any number of generations. (**B**) We collected data on estimates of reproductive isolation between populations that have been evolving in the same environment (within environments, in blue, A1 × A2 and B1 × B2) or in different environments (between environments, in orange, A1 × B1, A1 × B2, A2 × B1 and A2 × B2). If estimates of reproductive isolation are greater between environments than within environments, this is evidence for ecological speciation, whereas if they are equal, this provides evidence for mutation order speciation. (**C**) The dataset is predominantly comprised of data from invertebrates that are best suited for experimental evolution studies, like *Drosophila melanogaster* (C1), *Ceratitis capitata* (C2), and *Tetranychus urticae* (C3), though some vertebrate translocation experiments, like those done in *Poecilia reticulata* (C4) also fit the criteria. (**D**) The phylogeny of all the species included in the dataset, from yeast to nematodes to vertebrates. Silhouettes are all public domain obtained from phylopic.org. Photo credits: (C1) Alexis, iNaturalist, CC-by-4.0 license; (C2) Jari Segreto, Flickr, CC-by-2.0 license; (C3) Gillis San Martin, Flickr, CC-by-2.0 license; (C4) Amy E. Deacon, Hideyasu Shimadzu, Maria Dornelas, Indar W. Ramnarine and Anne E. Magurran, CC-by-4.0 license.

### Database of experimental speciation experiments

We performed a literature search for experimental speciation experiments published in the thirty years after the publication of Rice and Hostert (*16*) (see Methods). Rice and Hostert (*16*) stimulated several new experimental evolution studies testing causal mechanisms underlying the evolution of reproductive isolation (*31, 32*), now enabling our formal meta-analysis. Though multiple mechanisms have been investigated using experimental evolution studies, including the roles of genetic bottlenecks and sexual conflict, we focus on if and how divergent selection could increase the rate of evolution of reproductive isolation. We compiled a dataset of 1723 effect sizes from 34 studies on 18 species that imposed divergent selection and estimated reproductive isolation within and between selective regimes (Methods). Estimates of reproductive isolation *within* environments allow us to quantify the role of mutation-order speciation; that is, populations that evolve reproductive isolation in response to the same environment. Estimates of reproductive isolation *between* populations subject to different selection regimes allow us to quantify the role of ecological speciation (Figure 1B).

As effect sizes, we used the metric of Sobel and Chen (*35*), which quantifies reproductive isolation between populations as

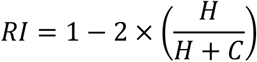

where H is the number or frequency of heterospecific/heterotypic matings, and C is the number or frequency of conspecific/homotypic matings. This framework can be readily applied for both pre-mating and post-mating isolation where H can be substituted for immigrant or hybrid fitness and C for resident fitness. This equation can be expanded to include any isolating mechanism separately and place it on the same scale of −1 to +1, where +1 is complete reproductive isolation, −1 is complete gene flow between the two populations (complete disassortative mating, for example), and 0 is random gene flow between populations. As this metric is equivalent to Pearson’s correlation coefficient, we used Fisher’s z-transformation to normalise it (*zRI*) and used 1 / n−3 as an estimate for the sampling variance for each effect size, where n is the sample size (see Supplementary Materials for sensitivity analyses using a different estimate for the sampling variance).

Unsurprisingly, invertebrates dominate the dataset (16 / 18 species, Figure 1C and D, Table S3), with *Drosophila* species making up the majority (35.3%) of the experiments, as n Rice and Hostert (*16*). Here, we fit a grand meta-analytic model including all the variables subsequently listed. We classified each estimate of reproductive isolation as a pre-mating or post-mating barrier. Pre-mating isolation barriers were reported more commonly than post-mating barriers, representing 91.7% of the total effect sizes, and sexual isolation being the most common pre-mating barrier estimated (see Supplementary Information for a detailed breakdown of the effect sizes. We also collected data on: 1) how many generations the experiment had been running when reproductive isolation was estimated (median = 43 generations, range = 8–1589); 2) founding population size for each replicate (median = 280, range = 1–5000); 3) whether or not populations experienced a common garden generation to minimise environmental effects on the estimate (*36*); 4) whether estimates were at the replicate population level or if estimates involved more than two populations (see Methods); and, 5) if our estimate of reproductive isolation was calculated from means and standard errors from papers where raw data were not available.

Finally, we collected information on whether reproductive isolation was measured between populations evolving in the same environment, or whether the two populations experienced divergent selection imposed by different environments. If divergent selection and ecological differentiation increase the rate of evolution of reproductive isolation, reproductive isolation would be expected to be greater between populations evolving in different environments compared to populations evolving in the same environment.

### Divergent selection increases reproductive isolation

Populations that evolved in different environments exhibited greater reproductive isolation than populations that had evolved in the same environment (difference estimate = 0.074, 95% CIs = [0.040, 0.106], pMCMC < 0.001, Figure 2A). This result aligns with the predictions made by ecological speciation theory (*18*), and with the initial conclusions reached by Rice and Hostert (*16*) three decades ago. Adaptation to different environments likely accelerates the build-up of genetic differences between populations, which increases the likelihood that alleles that contribute to reproductive isolation are linked with selected alleles (*37*). Alternatively, selected alleles pleiotropically contribute to reproductive isolation (*38*). These results suggest that the evolution of reproductive isolation by divergent selection may be common and detectable across a range of taxa and environments.

**Figure 2.**
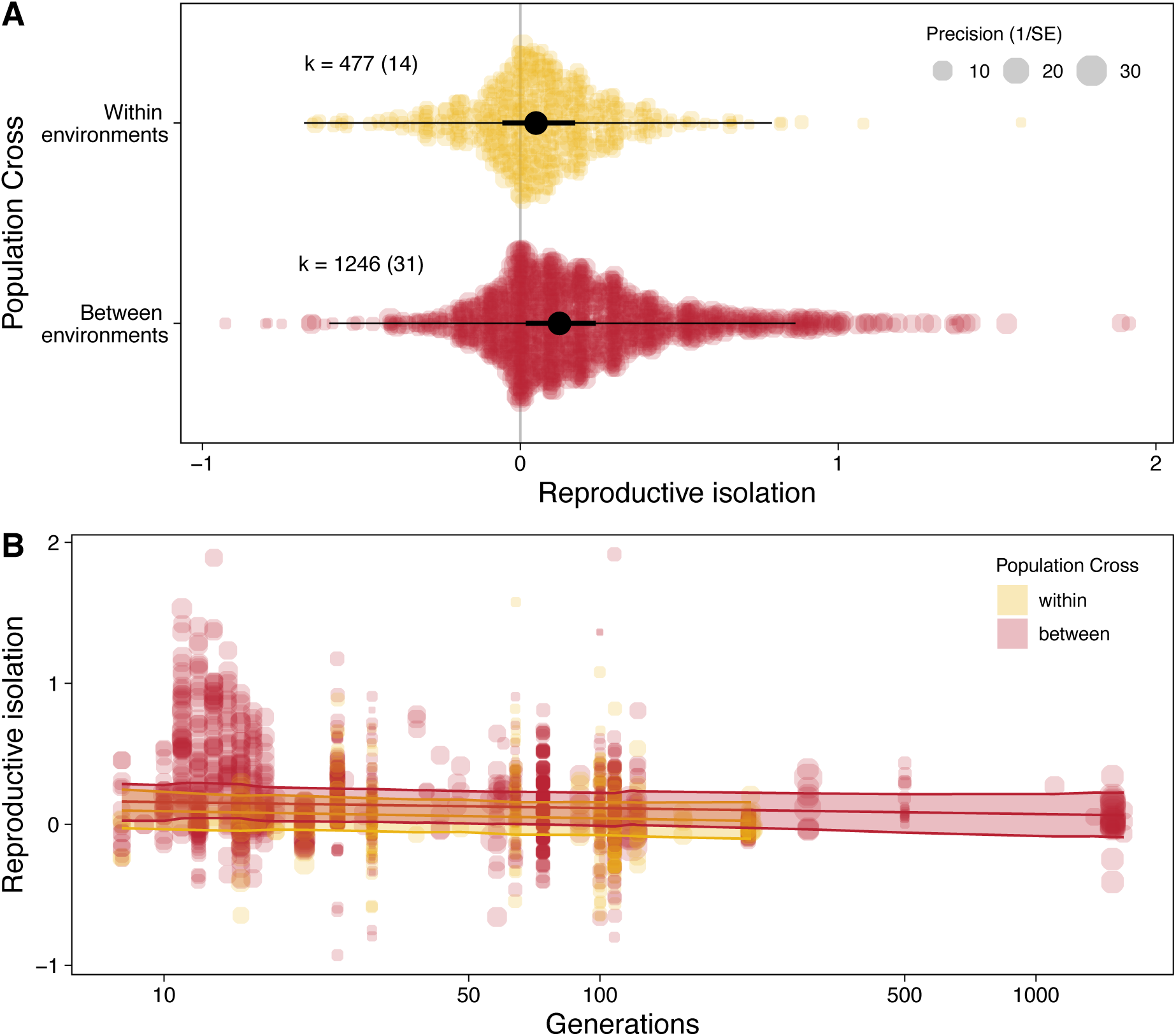
The evolution of reproductive isolation as a byproduct of divergent selection. (**A**) Reproductive isolation is greater for populations have been evolving in different environments (between environments, red) compared to populations that have been evolving in the same environment (within environments, yellow). k indicates the number of effects sizes extracted with the number of studies in brackets. Thick black lines are the 95% confidence intervals and the thin black lines are the prediction intervals. (**B**) However, the magnitude of reproductive isolation is independent of the number of generations of the experimental evolution study. Reproductive isolation is z-transformed. Each point is an effect size. The size of the circle indicates the precision (1 / SE) of the effect size, as shown in the legend in A. The shaded area indicates the 95% confidence interval for the line

Diet was the predominant type of environment and form of divergent selection in these experimental evolution studies (55.9% of studies, Supplementary Information). While diet may select on a range of phenotypic traits, most studies actively manipulated a single dimension of divergent selection. The two exceptions were Castillo et al. (*39*) who manipulated both diet and access to food, and Rundle (*40*), who used ancestral and novel environments that differed in diet, temperature, photoperiod, and feeding schedule. While our data suggest that a single environmental dimension of divergent selection is sufficient to promote the evolution of reproductive isolation, multidimensional or multifarious selection has been suggested to accelerate adaptation (*16, 41*). However, there is currently little direct empirical data to support the claims that multifarious selection can accelerate the evolution of reproductive isolation (*42, 43*). Manipulating multiple environmental axes to generate multifarious divergent selection should be a high priority in future experimental speciation studies, as changes in ecology are likely to be multidimensional (*32*).

### Reproductive isolation does not increase with time

A central tenet of speciation research is that reproductive isolation should increase with time as two populations or incipient species diverge (*1*). The shape of the relationship between reproductive isolation and time (or genetic distance) would depend on the mechanism underlying reproductive isolation. Intrinsic hybrid incompatibilities can accumulate at a rate faster than linear (*29*), and pre-zygotic isolation evolves more rapidly in sympatric than in allopatric *Drosophila* species (*27*). To understand the dynamics of reproductive isolation at the onset of speciation, we tested for the presence of an interaction between the population cross (within or between environments) and the number of generations of evolution. Our prediction was that the between-environment slope of how reproductive isolation develops with time would be greater than the within-environment slope, leading to the end result that divergent selection increases reproductive isolation (Figure 2A).

Surprisingly, we did not find evidence of an interaction (slope difference estimate = −0.001, 95% Cis = [−0.040, 0.040], pMCMC = 0.962); nor did we find an effect of the number of generations at all (slope estimate = −0.002, 95% CIs = [−0.032, 0.034], pMCMC = 0.92). Reproductive isolation, therefore, does not increase with time in experimental speciation studies (Figure 2B). One explanation for this unexpected result is that the environmental manipulations designed by experimenters were of such a large magnitude of selection that adaptation and concomitant reproductive isolation occurred rapidly and developed almost instantaneously. The major part of reproductive isolation would then be expected to have evolved in the earliest stages of the experiments. This hypothesis is difficult to test as reproductive isolation was usually measured once or twice during the course of a single experiment, and so the temporal dynamics of reproductive isolation may be obscured by the different experimental study designs. To circumvent this problem, we analysed a subset of our data containing those studies that measured reproductive isolation at three or more time points (N = 5 studies, Supplementary Information). The average slope was not significantly different from zero in this subset of studies either (slope estimate = −0.22, 95% CIs = [−0.99, 0.62], pMCMC = 0.47).

A second explanation is that the strength of selection was not of equal magnitude across studies. This would have resulted in different rates of adaptation and thus unequal probabilities of the evolution of reproductive isolation. Such between-study heterogeneity may have obscured a clear pattern between reproductive isolation and time. A third explanation could be that populations with smaller effective population sizes may have fixed alleles faster due to genetic drift, compared with larger populations. This would lead to forestalled adaptation and a cessation of the gradual build-up of reproductive isolation. To test this hypothesis, we examined the founding population size reported by each study. However, we found no significant relationship between founding population size and the development of reproductive isolation (slope estimate = 0.005, 95% CIs = [−0.009, 0.024], pMCMC = 0.524).

### Phenotypic plasticity accelerates the emergence of reproductive isolation

A fourth explanation is that developmental plasticity induced by the divergent environments causes reproductive isolation itself within a single generation (*44–46*). We tested for any role of the developmental environment on reproductive isolation by leveraging the variation across studies in their use of a common garden generation. A common garden generation involves rearing all populations in the same environment to remove effects of plasticity prior to assessing any evolutionary change that has occurred in the populations (*36*). We asked if reproductive isolation was greater when a common garden was not used to purge the environment effects on phenotypes, and thus plasticity as well as any genetic differences between populations contributed to reproductive isolation. We found that plasticity likely plays an important role in promoting the early emergence of reproductive isolation (Figure 3). Specifically, populations that experienced a common garden generation prior to the estimation of reproductive isolation exhibited significantly lower estimates of reproductive isolation than populations not exposed to a common garden environment (difference estimate = −0.109, 95% CIs = [−0.199, −0.021], pMCMC = 0.016).

**Figure 3.**
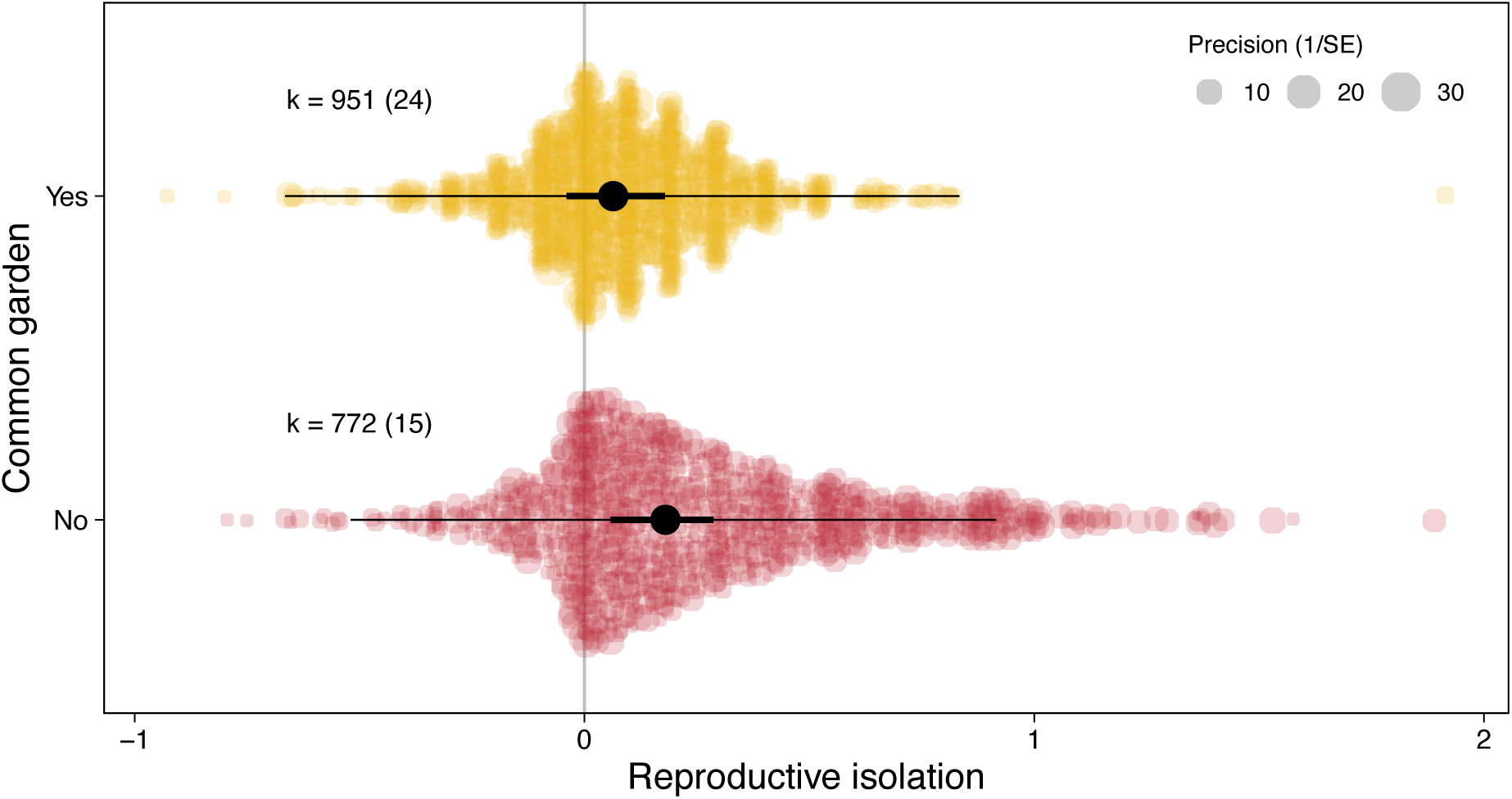
Plasticity increases reproductive isolation. Populations that were reared in a common garden environment had lower estimates of reproductive isolation than populations that did not pass through a common garden environment. Reproductive isolation is z-transformed. k indicates the number of effects sizes extracted with the number of studies in brackets. Thick black lines are the 95% confidence intervals and the thin black lines are the prediction intervals.

Developmental plasticity and learning are two ways by which phenotypic plasticity can promote reproductive isolation. When reproductive isolation is linked to morphological traits, developmental plasticity could increase the phenotypic differentiation between populations over and above any genetic differentiation that may have evolved, thus increasing the degree to which the populations are reproductively isolated from one another. For example, diet, a common environment used in the studies here, can induce morphological shifts in body size that in turn can result in reproductive isolation between populations that developed in different environments (*47*). In addition, environmentally induced differences in other morphological traits or chemical signals could accentuate underlying patterns of assortative mating that already exist within populations (*48–51*). owever, without a correlation between parental and offspring environments, environmentally induced assortative mating will have little effect on speciation dynamics (*45*). Learning is a special form of plasticity that can immediately impact the extent of gene flow between populations (*52, 53*). Offspring can imprint on their natal habitat (*54, 55*), which, in combination with environmentally induced assortative mating, could promote reproductive isolation. Offspring can also imprint on their parents’ phenotypes (*56–58*), or in the case of brood parasitic birds, their host species (*59, 60*). Although environmental factors can promote reproductive isolation through various forms of plasticity, such environmentally induced reproductive isolation is likely to be fragile and collapse as soon as the environment changes. Breakdown of ecological differences between recently diverged populations could then result in incomplete speciation (*42*) or speciation reversal (*61*), with species persisting only when intrinsic post-mating isolating barriers have subsequently evolved (*13*).

### Evolution of isolating barriers mirrors macroevolutionary patterns

Our study focuses on the onset of speciation, but are these findings also relevant to the end of speciation? The link between the microevolution of reproductive isolation and macroevolutionary speciation rates has been subject to much recent discussion, and is far from resolved (*5, 49, 62*). Comparative analyses suggest that pre-mating isolating barriers likely evolve earlier in the speciation process and are usually initially stronger than post-mating barriers (*63, 64*), especially when taxa are sympatric, suggestive of reinforcement (27). Consistent with this, we found that in experimental speciation studies, pre-mating reproductive isolation is stronger than post-mating reproductive isolation (difference estimate = 0.149, 95% CIs = [0.072, 0.221], pMCMC < 0.001, Figure 4). In many cases, however, post-mating isolating barriers evolve just as fast, if not faster, than pre-mating barriers at least in some taxa (*51, 65*). Our meta-analysis on experimental speciation studies has highlighted the dearth in post-mating estimates of reproductive isolation as most studies have focused on the consequences of mating (e.g. decreases in longevity and fecundity). There are only a handful of estimates of hybrid fitness traits which are often measured at the latter end of the speciation continuum. A consensus of fitness measures that can be measured both in the laboratory and in the field is therefore clearly needed to align studies that focus on the onset of speciation, and its completion.

**Figure 4.**
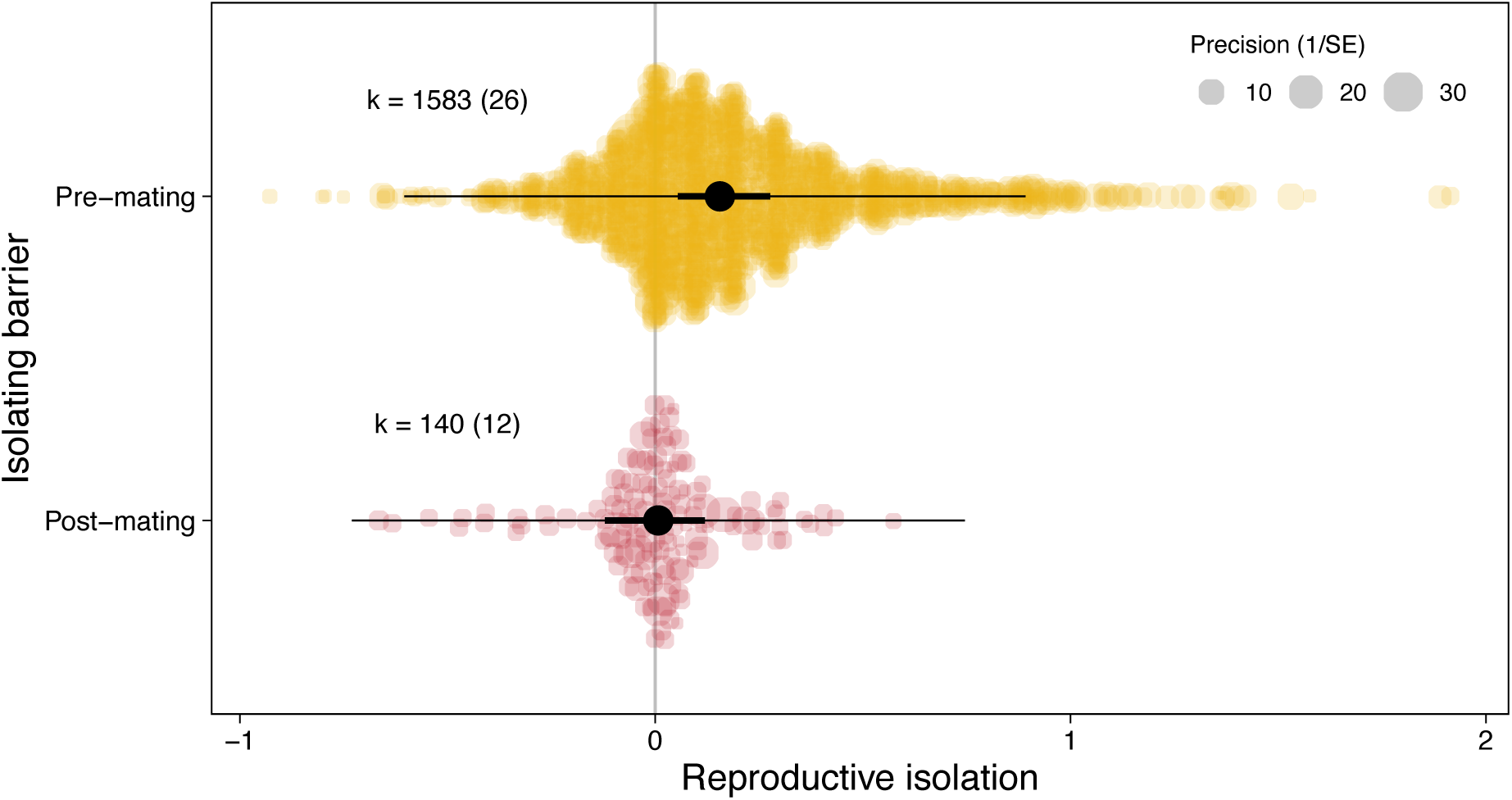
Pre-mating isolating barriers are more likely to evolve than post-mating isolating barriers in the early stages of speciation. The majority of effects sizes come from pre-mating isolating barriers (sexual isolation, habitat choice), with fewer estimates of post-mating barriers (hybrid fitness, longevity). Reproductive isolation is z-transformed. k indicates the number of effects sizes extracted with the number of studies in brackets. Thick black lines are the 95% confidence intervals and the thin black lines are the prediction intervals.

### Conclusions

Our results show that experimental speciation studies causally support the ecological speciation model for the evolution of reproductive isolation. Our meta-analysis provides quantitative evidence that: (i) divergent selection can promote the rapid evolution of reproductive isolation (Figure 2A); (ii) that phenotypic plasticity contributes to reproductive isolation (Figure 3); and (iii) that pre-mating isolating barriers evolve faster that post-mating barriers (Figure 4). Our study also shows that the evolutionary dynamics of reproductive isolation in the early stages of speciation do not necessarily match the dynamics further along the speciation continuum (Figure 2B). While recent work has suggested that ecological speciation may not be the most common mechanism by which species originate (*23*), our results suggest that divergent selection can rapidly form populations that display significant reproductive isolation, consistent with ecological speciation theory. Further, we show that plasticity may be key in initiating reproductive isolation, but its role in sustaining reproductive isolation may depend on the evolution of plasticity itself (*66*).

A few model organisms have dominated ecological speciation research: anole lizards, stickleback fish (*Gasterosteus aculeatus*), cichlid fish, *Timema* walking sticks, *Rhagoletis* flies, and cynipid gall wasps. These model systems have all provided key insights into the process of speciation, but seldom measure the evolution of reproductive isolation in real time, across generational scales. The strength of the experimental speciation approach is that it has revealed the potential and importance of ecological speciation more generally and across a wider range of taxa. Future experimental speciation studies should broaden both the species used and the dimensionality of selection, whilst measuring multiple isolating barriers (both pre- and post-mating) at multiple timepoints to further our causal understanding of the evolution of reproductive isolation in the early stages of speciation.

## Acknowledgements

We thank M. Björklund, A. Chippindale, S. Magalhães, N. G. Prasad, H. Rundle, V. Shenoi, and H. Thyagarajan for kindly provided requested data, and the permission to upload the data with this manuscript. We also thank A. Comeault, S. Nilén, D. Parker, I. Prates, and M. Tsuboi for comments.

## Author contributions

B.J.M.J and E.I.S conceived the study, B.J.M.J collected the data, B.J.M.J and P.A.D analysed the data, B.J.M.J wrote the first draft, all authors contributed to the final version.

## Data and materials availability

All data and code are available in the manuscript or the supplementary materials.

## Materials and Methods

### 1. Overview

To quantitatively synthesise studies on the experimental evolution of reproductive isolation, we conducted a systematic literature search and meta-analysis. This involved five main steps: i) defining study eligibility criteria; ii) building and validating a search string to find studies meeting these criteria; iii) conducting our search in Web of Science and Scopus and screening studies; iv) calculating effect sizes from our final sample of studies; and, v) building statistical models to estimate mean effect sizes and to explore the effects of generation time and study design on estimates of reproductive isolation.

**Figure S1.**
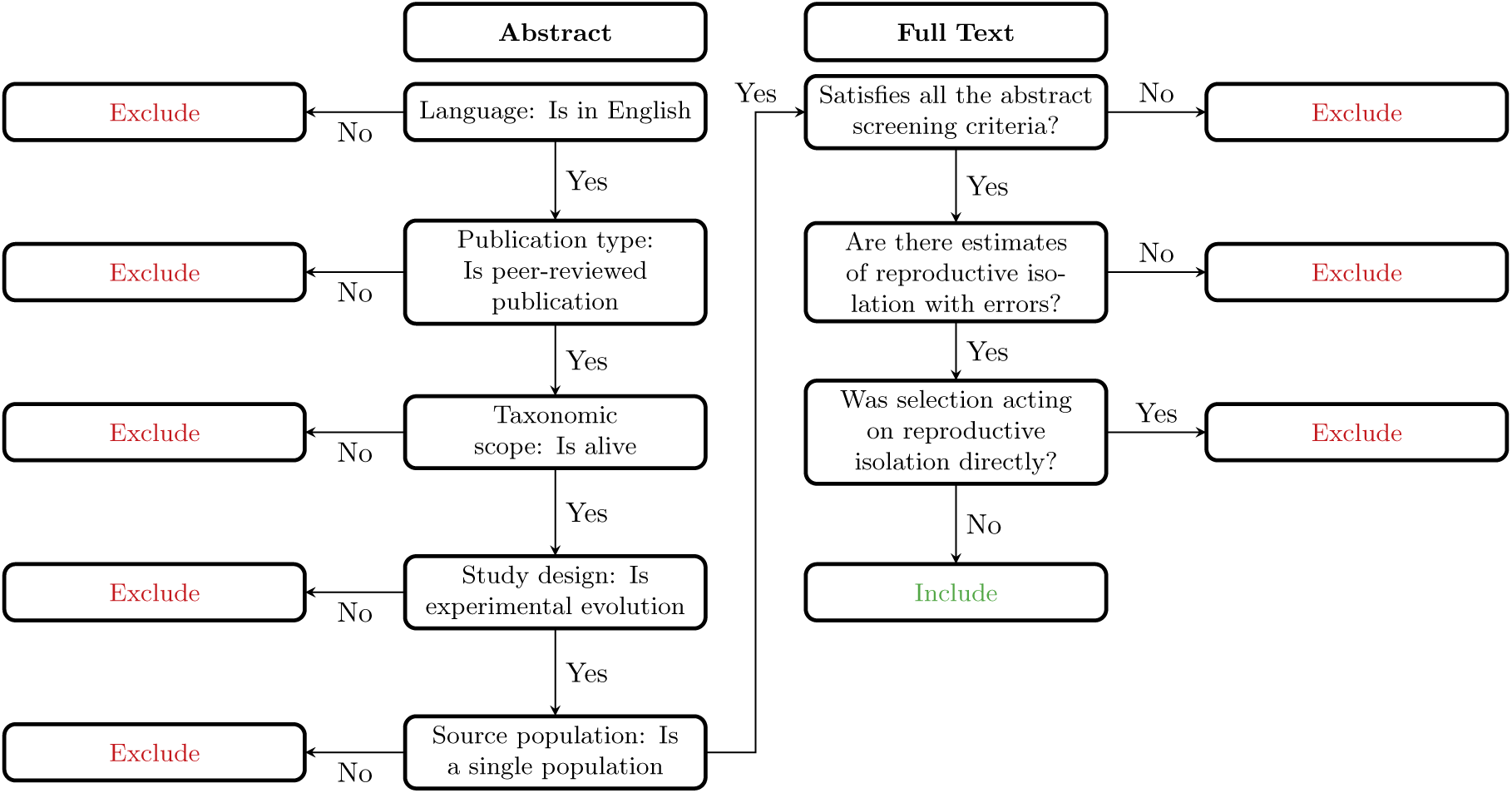
Decision tree for title and abstract screening, and full text screening.

### 2. Eligibility criteria

To be included in our dataset, studies had to meet the following criteria: 1) the study had to be in English; 2) the study is a peer-reviewed research paper; 3) the study needed to include data from alive organisms (we were taxon unspecific); 4) the study used experimental evolution with a founding population from a single genetic source (i.e. not from multiple wild populations that may or may not have experienced different selective pressures with its own genetic architectures that may or may not bias the path of evolution and subsequent reproductive isolation); 5) data had to be available, either as raw data or as means with associated standard errors/deviations and sample sizes; and, 6) selection did not directly act on reproductive isolation itself (e.g. studies were excluded if selection acted on assortative mating). Our PICO components were: Population, experimentally-evolved living organisms; Intervention, experimental evolution in different environments and with measures of reproductive isolation between populations; Comparison, estimates of reproductive isolation between populations evolving in the same environment compared to populations evolving in different environments; Outcome, estimate of reproductive isolation acting via any isolating barrier. From this, we constructed two decision trees, one for title and abstract screening and one for full text screening (Figure S1).

**Table S1.**
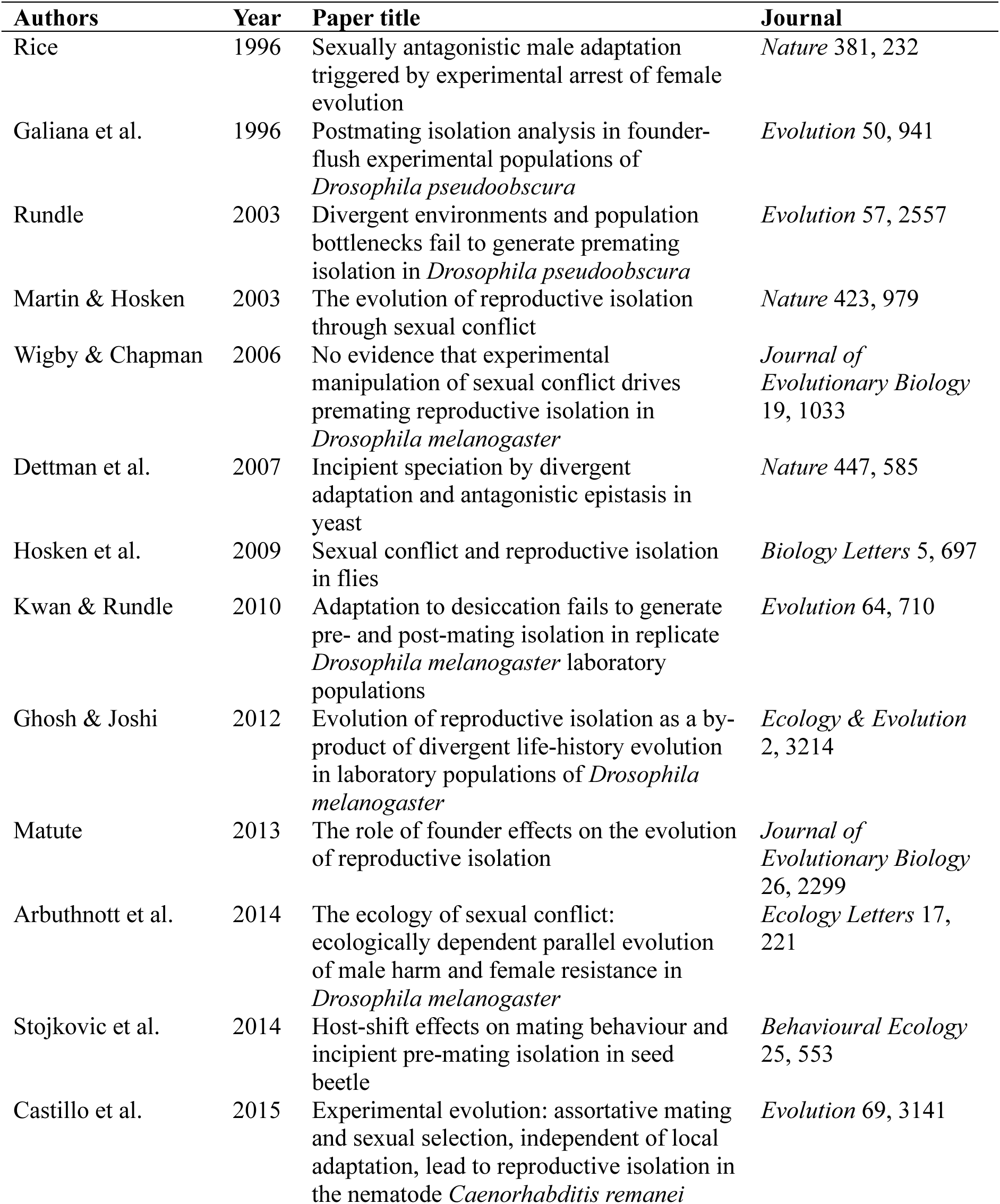
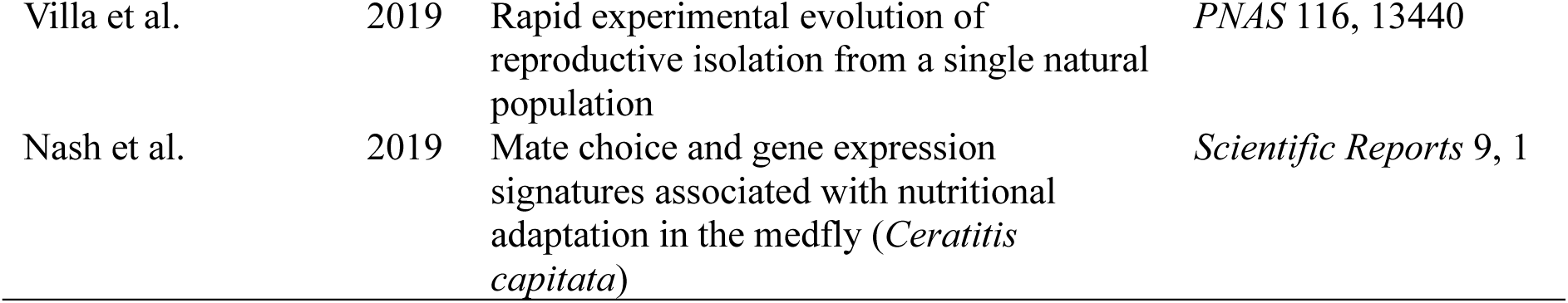
List of benchmarking papers used for our meta-analysis search. All papers were found in our literature search.

| Authors | Year | Paper title | Journal |
| --- | --- | --- | --- |
| Rice | 1996 | Sexually antagonistic male adaptation triggered by experimental arrest of female evolution | *Nature* 381, 232 |
| Galiana et al. | 1996 | Postmating isolation analysis in founder-flush experimental populations of *Drosophila pseudoobscura* | *Evolution* 50, 941 |
| Rundle | 2003 | Divergent environments and population bottlenecks fail to generate premating isolation in *Drosophila pseudoobscura* | *Evolution* 57, 2557 |
| Martin & Hosken | 2003 | The evolution of reproductive isolation through sexual conflict | *Nature* 423, 979 |
| Wigby & Chapman | 2006 | No evidence that experimental manipulation of sexual conflict drives premating reproductive isolation in *Drosophila melanogaster* | *Journal of Evolutionary Biology* 19, 1033 |
| Dettman et al. | 2007 | Incipient speciation by divergent adaptation and antagonistic epistasis in yeast | *Nature* 447, 585 |
| Hosken et al. | 2009 | Sexual conflict and reproductive isolation in flies | *Biology Letters* 5, 697 |
| Kwan & Rundle | 2010 | Adaptation to desiccation fails to generate pre- and post-mating isolation in replicate *Drosophila melanogaster* laboratory populations | *Evolution* 64, 710 |
| Ghosh & Joshi | 2012 | Evolution of reproductive isolation as a by-product of divergent life-history evolution in laboratory populations of *Drosophila melanogaster* | *Ecology & Evolution* 2, 3214 |
| Matute | 2013 | The role of founder effects on the evolution of reproductive isolation | *Journal of Evolutionary Biology* 26, 2299 |
| Arbuthnott et al. | 2014 | The ecology of sexual conflict: ecologically dependent parallel evolution of male harm and female resistance in *Drosophila melanogaster* | *Ecology Letters* 17, 221 |
| Stojkovic et al. | 2014 | Host-shift effects on mating behaviour and incipient pre-mating isolation in seed beetle | *Behavioural Ecology* 25, 553 |
| Castillo et al. | 2015 | Experimental evolution: assortative mating and sexual selection, independent of local adaptation, lead to reproductive isolation in the nematode *Caenorhabditis remanei* | *Evolution* 69, 3141 |
| Villa et al. | 2019 | Rapid experimental evolution of reproductive isolation from a single natural population | *PNAS* 116, 13440 |
| Nash et al. | 2019 | Mate choice and gene expression signatures associated with nutritional adaptation in the medfly (*Ceratitis capitata*) | *Scientific Reports* 9, 1 |

### 3. Search string development and validation

To build our search strings, we chose 15 benchmark studies that we wanted our literature search to return (Table S1). These studies were chosen for their importance in testing the idea of divergent selection, sexual conflict, and genetic bottlenecks in generating reproductive isolation using experimental evolution. The titles and abstracts from these studies were used to construct a word cloud (Figure S2), which enabled us to identify common terms and synonyms used across studies. We wanted our search to return between 1000 and 3000 studies. The final search strings (separated by **AND**) were:

*“divergent selection” AND hybrid* **AND** *“experimental evolution” AND “ecological speciation”* **AND** *“experimental evolution” AND speciation* **AND** *“experimental evolution” AND “reproductive isolat*”* **AND** *“hybrid inviab*” OR “hybrid steril*” AND “experimental evolution”* **AND** *(prezygotic OR postzygotic OR pre-zygotic OR post-zygotic) isolation AND “experimental evolution”* **AND** *“experimental evolution” AND “sexual isolation”* **AND** *“host plant” AND hybrid AND isolati\** **AND** *“experimental evolution” AND adapt* AND divergen* AND speciation* **AND** *bottleneck AND “reproductive isolat*” **AND** bottleneck AND speciation AND “experimental evolution”* **AND** *“founder flush” OR “founder-flush”*.

We validated the search string, before title and abstract screening all articles, by calculating its miss rate (the % of the benchmark papers missing) and hit rate (the % of papers that pass to full text screening out of random 100 papers). The miss rate was 13% (2 / 15 benchmark studies were missing) and the hit rate was 7% (of 100 random papers, 7 passed to full text screening), which are typical in the field. Note, that we used our title and abstract decision tree (Figure S1) to calculate the hit rate.

**Figure S2.**
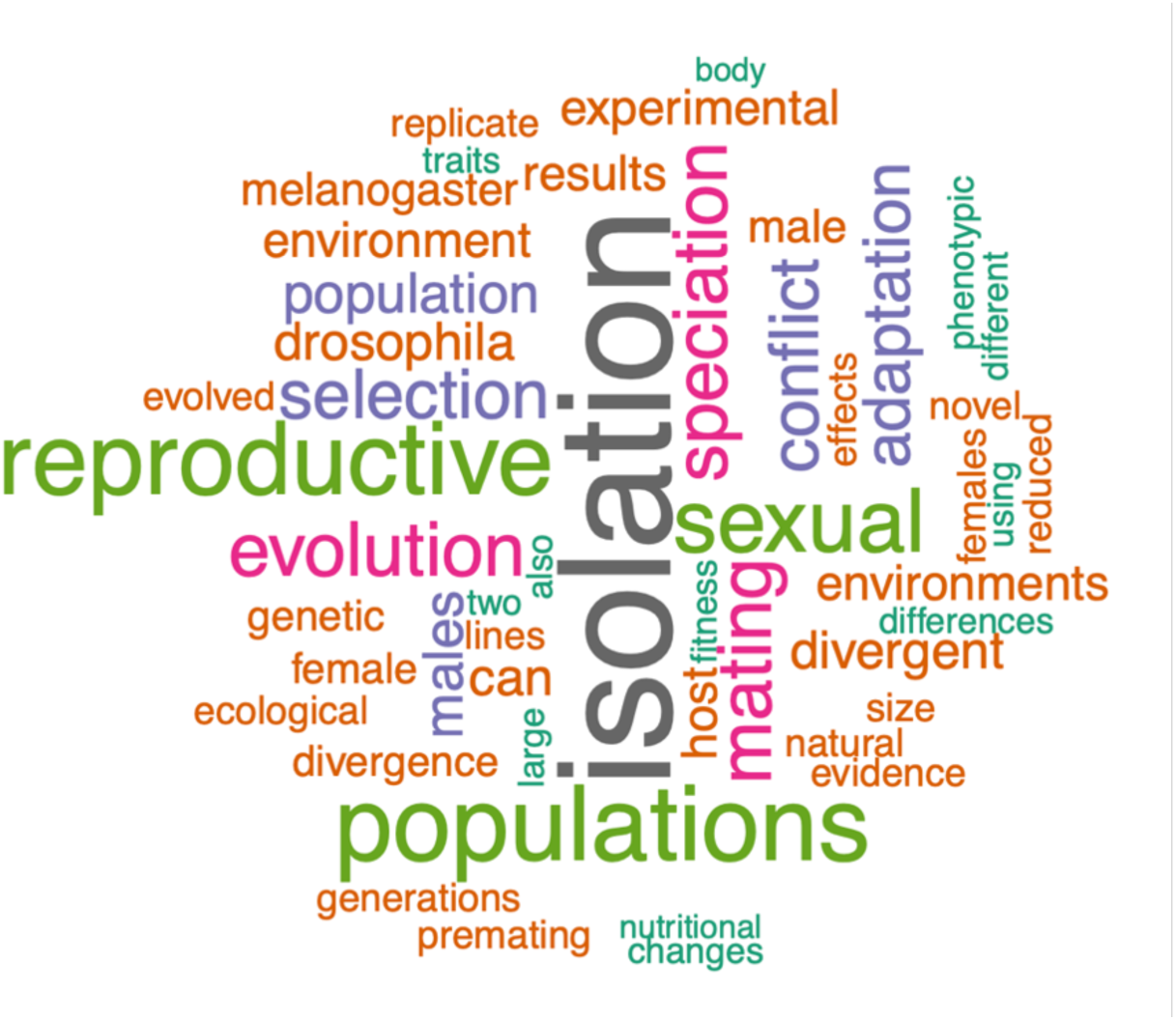
Word cloud formed from the title and abstracts of the 15 benchmarking papers (Table S1). This guided our construction of search strings.

### 4. Searches and screening

We performed two topic searches using Web of Science and Scopus, adapting out search string for each. The first search was done in March 2021 (from Lund University, Sweden) to search for relevant studies, and the second in June 2023 (from Bangor University, UK) to account for studies published since our first search. The databases covered by Web of Science in each institution are reported in Table S2. Only studies published in 1994 and beyond (after Rice and Hostert (*16*), published in December 1993) were considered.

**Table S2.**
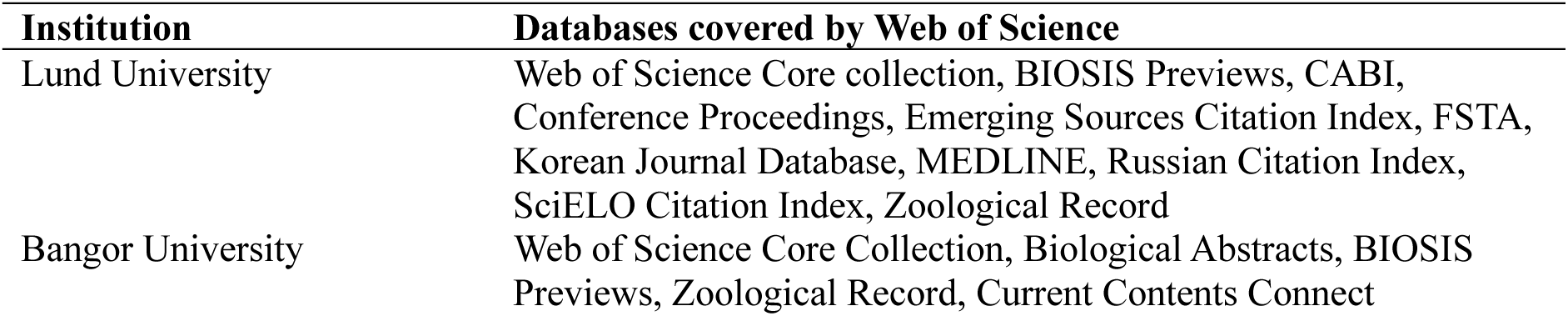
Databases covered by Web of Science at the two universities where searches were performed. The Scopus search only searched the Scopus database.

In total, our literature search returned 2715 studies (Figure S3). After removing 1008 duplicates, we screened the titles and abstracts for 1707 studies in Rayyan (*67*) following our decision tree. We selected 82 studies for full text screening to which we added nine studies from other sources, which mainly included citations in screened papers (*68–76*), and five studies cited in Edelaar (*77*), a review published in 2022 (*78–82*). Of these 96 studies, 48 were included, with 34 studies eligible for the meta-analysis concerned with the role of divergent selection in generating reproductive isolation (Table 3).

**Figure S3.**
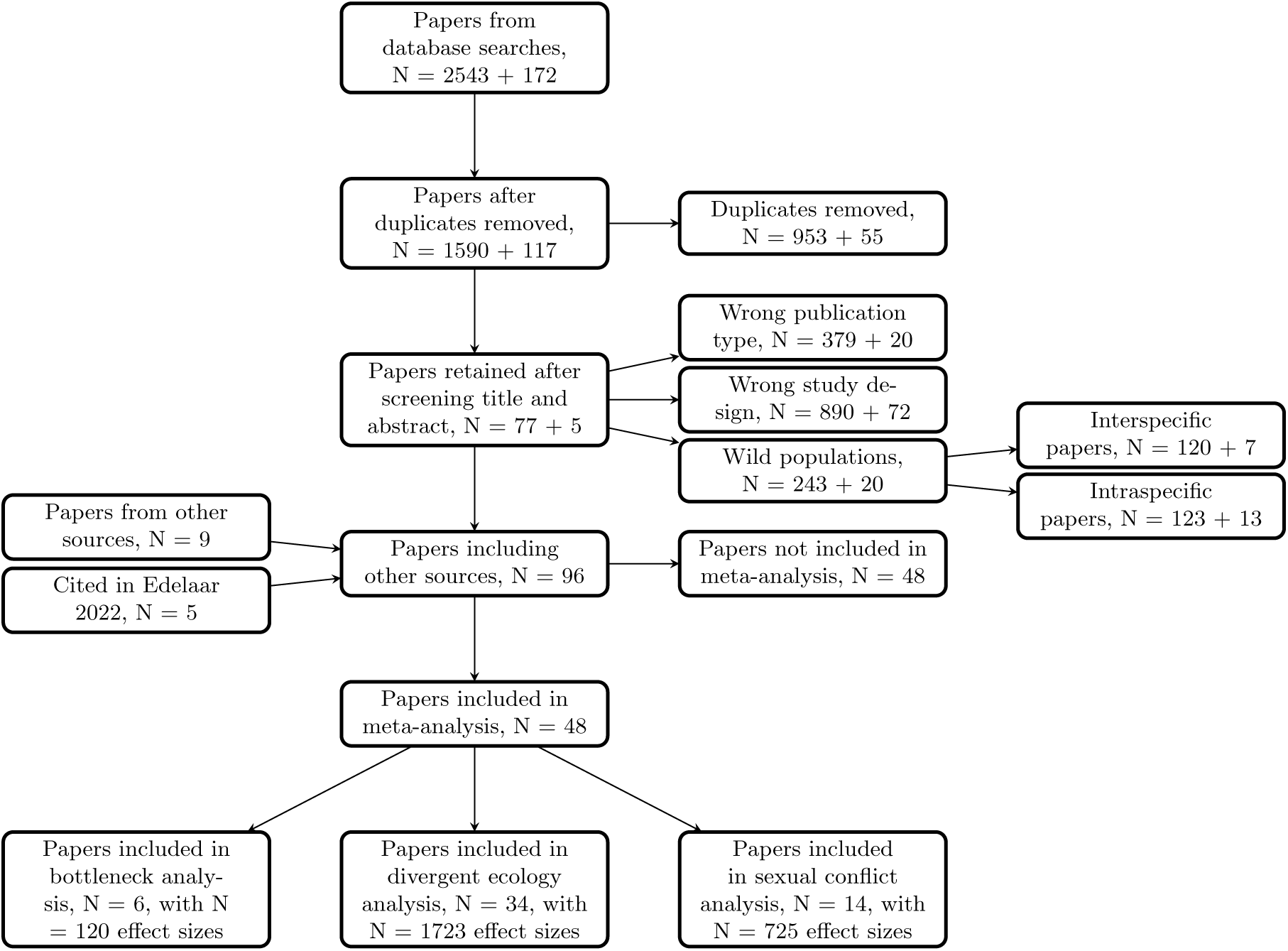
PRISMA flow chart. The two numbers in each block are from the two searches (the first in March 2021 at Lund University and the other in June 2023 at Bangor University. This manuscript covers only the results extracted from the middle block at the bottom “*Papers included in the divergent ecology analysis*”. The blocks either side are included for completeness.

### 5. Data extraction and effect size calculation

All data manipulation and analysis was done in R (*111*). As an effect size, we used the metric of Sobel and Chen (*35*), which quantifies reproductive isolation between populations as

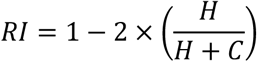

where H is the number or frequency of heterospecific/heterotypic matings, and C is the number or frequency of conspecific/homotypic matings. This framework can be readily applied for both pre-zygotic and post-zygotic isolation where H can be substituted for immigrant or hybrid fitness and C for resident fitness. This equation can be expanded to include any isolating mechanism separately and place it on the same scale of −1 to +1, where +1 is complete reproductive isolation, −1 is complete gene flow between the two populations (complete disassortative mating, for example), and 0 is random gene flow between populations. As this metric is equivalent to the Pearson’s correlation coefficient, we used Fisher’s z-transformation to normalise it (*zRI*) and used as an estimate for 1 / n−3 the sampling variance for each effect size, where n is the sample size.

Reproductive isolation can be viewed as an emergent property of the interaction between two populations. We therefore sought to include all estimates of reproductive isolation at the population level. For example, if a study had two replicate populations (1 and 2) evolving in two different environments (A and B), we would extract two within-treatment estimates of reproductive isolation (A1 × A2, and B1 × B2), and four between-treatment estimates of reproductive isolation (A1 × B1, A1 × B2, A2 × B1, and A2 × B2). As the same populations contribute to different estimates of reproductive isolation from the same paper, we accounted for this by calculating within-research group variance (co)variance matrices for each specific population used. To do this, we used the *make_VCV_matrix* function from the “metaAidR” package (*112*). In some cases, population-level estimates of reproductive isolation were not possible. This may be because the raw data were not made available to us, or because population-level estimates were underpowered, as many papers pooled individuals from different populations to assess the broad effects of different environments on the evolution of reproductive isolation. Effect sizes that were estimates at the population-population level are coded as “1” in the “pop_level_data” column of (1283 / 1723 effect sizes), with “0”s marking estimates of reproductive isolation where pooling of replicate populations occurred at any stage.

To calculate effect sizes, we preferred to use the raw data provided in the supplementary information (1542 / 1723 effect sizes). Raw data were reanalysed using the “brms” R package (*113, 114*) using binomial, beta, or bernoulli distributions with logit link functions for mate choice data, a gaussian distribution for relative fitness data, or a Gamma distribution with an exponential link function. For papers that did not provide supplementary data, we emailed the corresponding author to request it (98 / 1723 effect sizes). In cases where authors did not respond, we extracted means and errors from figures using the R package “metaDigitise” (*115*), randomly generating data with the same parameters, and estimated RI from these (83 / 1723 effect sizes). Effect sizes calculated this way are coded as “1” in the “backtransformed” column of the data.

#### 5.1 Meta data

We collected meta-data associated with our estimates of reproductive isolation. First, we grouped each estimate of reproductive isolation as to whether the isolating barrier operates “pre-mating” (1581 / 1723 effect sizes) or “post-mating” (142 / 1723 effect sizes). Pre-mating barriers operate before mating and fertilisation, and in our dataset include habitat selection (habitat choice or oviposition choice by females), with 184 / 1581 effect sizes, and, the most common form, sexual isolation (mate choice, latency to mate, and mating duration) with 1397 / 1581 effect sizes. Post-mating barriers operate after mating, and can either occur before or after fertilisation. Post-mating barriers can be intrinsic, where genetic incompatibilities exist between the two populations, or extrinsic, where the barrier operates via genetic differences between population but in conjunction with some aspect of the environment. In experimental speciation experiments, post-mating estimates of reproductive isolation are measured much less frequently, because they are more difficult to measure. Across the studies in our dataset, we classified post-mating isolation barriers into three broad categories: hybrid fitness (where hybrids are produced and fitness in any environment is measured, 34 / 142 effect sizes); fecundity (where the number of eggs or offspring is measured for hybrid crosses, 49 / 142 effect sizes); and female longevity (where female longevity is measured after mating or harassment from males from a different population, 59 / 142).

Second, we collected meta-data on whether reproductive isolation was estimated after every population had gone through a common garden generation or not. Common garden generations are used to remove effects of plasticity on trait values as all individuals experience the same developmental environment. Any differences in trait values or, in our case, reproductive isolation, could then be more confidently attributed to genetic changes in the population (*36*). In all papers, this information was readily available.

Third, we noted the generation of the experiment at which reproductive isolation was estimated. This was our estimate of divergence between populations. The median number of generations at which reproductive isolation was estimated in our dataset is 30 generations, with a range of eight and 1589 generations. We ln transformed the number of generations for inclusion in statistical models.

Lastly, initial population size may influence the rate at which populations evolve reproductive isolation. In a similar vein to genetic bottlenecks (papers related to which were uncovered by our literature search but are not the subject of this study), smaller populations may be more prone to genetic drift, where different alleles may become randomly fixed in different populations, with these genetic differences being responsible for reproductive isolation. Alternatively, larger populations may be more likely to exhibit reproductive isolation as adaptation to divergent selection and the build-up of genetic differences between populations may be more efficient than in smaller populations. We obtained this information from either the paper that reported estimates of reproductive isolation, or an earlier paper from the same research group that outlined the founding populations in more detail. We ln transformed the initial population size for inclusion in statistical models.

**Table S3.**
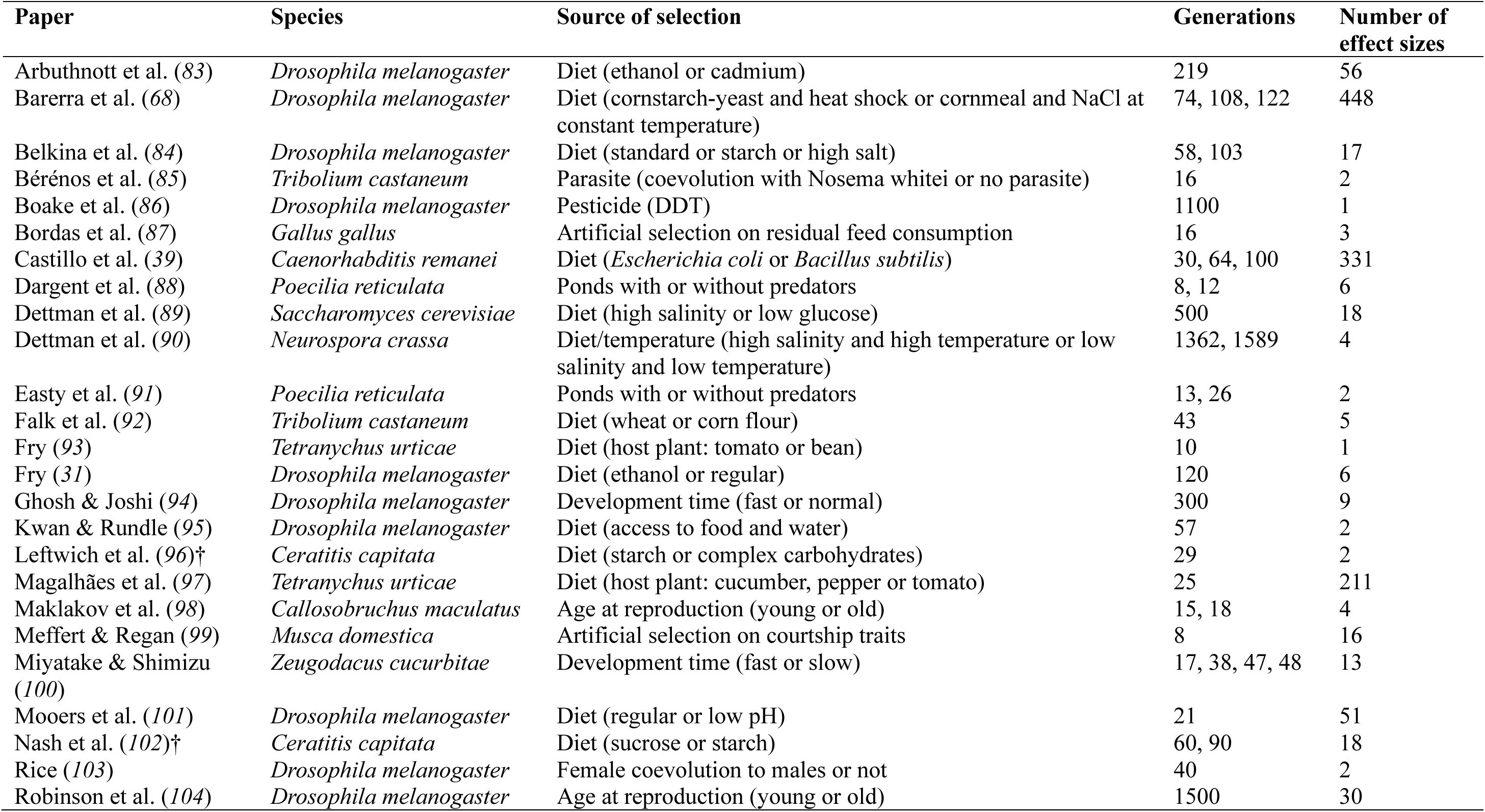
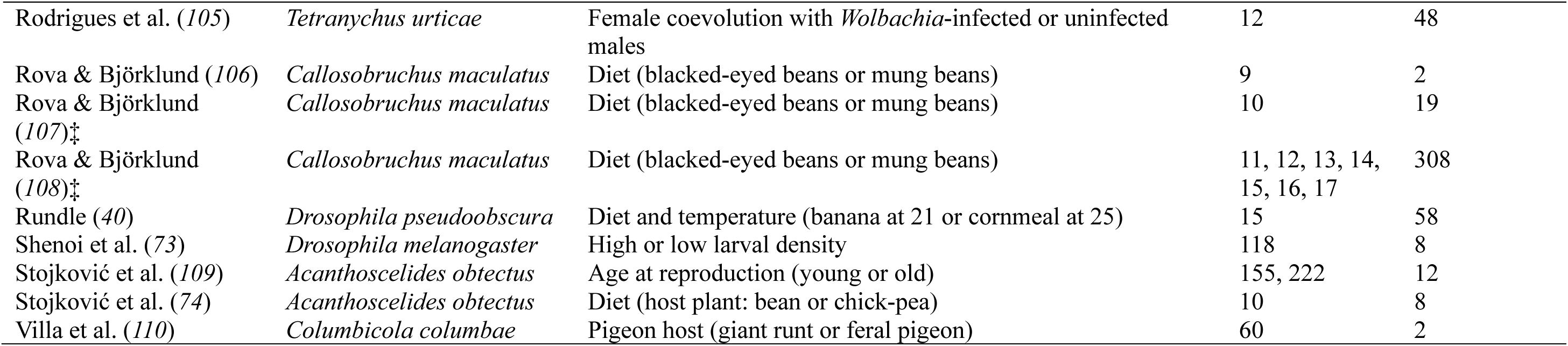
The list of papers included in the divergent selection dataset, with details of the species used, the selection imposed, when (in generations) reproductive isolation was estimated, and the amount of data each has contributed to the dataset. Two pairs of papers (indicated with the † and ‡ symbols) estimate reproductive isolation on the same populations.

### 6. Effect size summary

Invertebrates unsurprisingly dominate our dataset (Table S3). *Drosophila* (N_study_ = 12), *Callosobruchus maculatus* (N_study_ = 4), and *Tetranychus urticae* (N_study_ = 3) comprise the majority of the studies with other invertebrate species filling out the majority of the dataset. Two fungi species (*Saccharomyces cerevisiae* and *Neurospora crassa*) and two vertebrate species (the domesticated chicken *Gallus gallus* and the Trinidadian guppy *Poecilia reticulata*) make up the final species (Figure 1 and Figure S4).

### 7. Statistical analyses

#### 7.1 Generation of the phylogenetic tree

As multiple species are present in our dataset, we needed to control for phylogenetic signal that may be present in our dataset (i.e. closely related species may respond to divergent selection more similarly than more distantly related species). We used the “rotl” R package (*116*) to construct our cladogram, making use of the *tnrs_match_names* and *tol_induced_subtree* functions (Figure S4). We then used the Grafen method to compute the branch length using the “ape” package (*117*).

**Figure S4.**
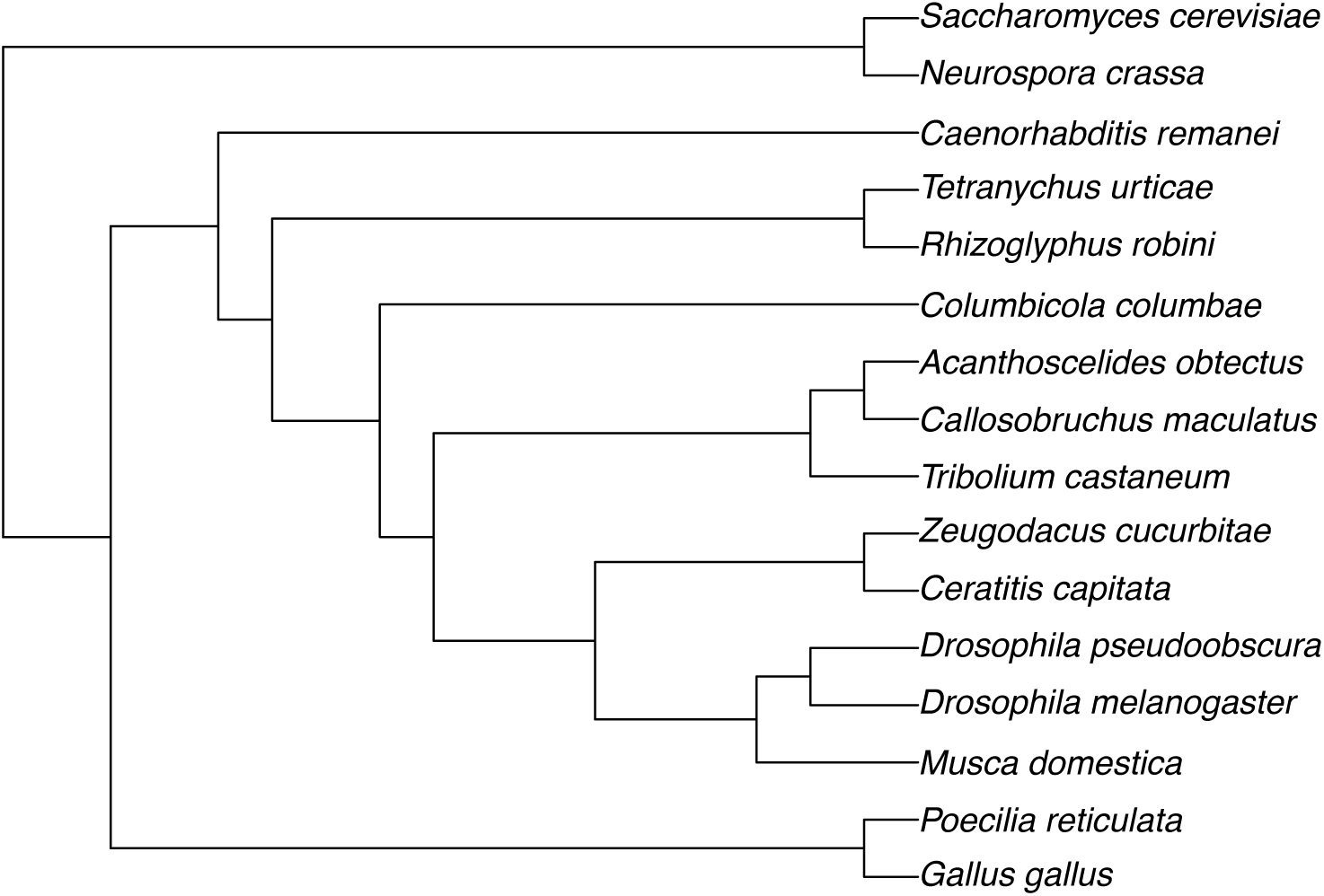
A cladogram showing the relationship between species used in the experimental speciation experiments that are included in our dataset.

#### 7.2 Publication bias and heterogeneity

We explored the effects of publication bias using five different methods: 1) funnel plots; 2) trim and fill analysis; 3) fail safe number; 4) Egger’s test; and 5) time lag analysis (*118*). Publication bias results from studies remaining unpublished (e.g. due to non-significant findings), which can skew the distribution of effect sizes in a meta-analysis. Models investigating publication bias and heterogeneity are included in the related code and were performed both in “MCMCglmm” (*119*) and “metafor” (*120*) in R (*111*). All priors and random effects are consistent between these models and the models used for the main analyses.

**Figure S5.**
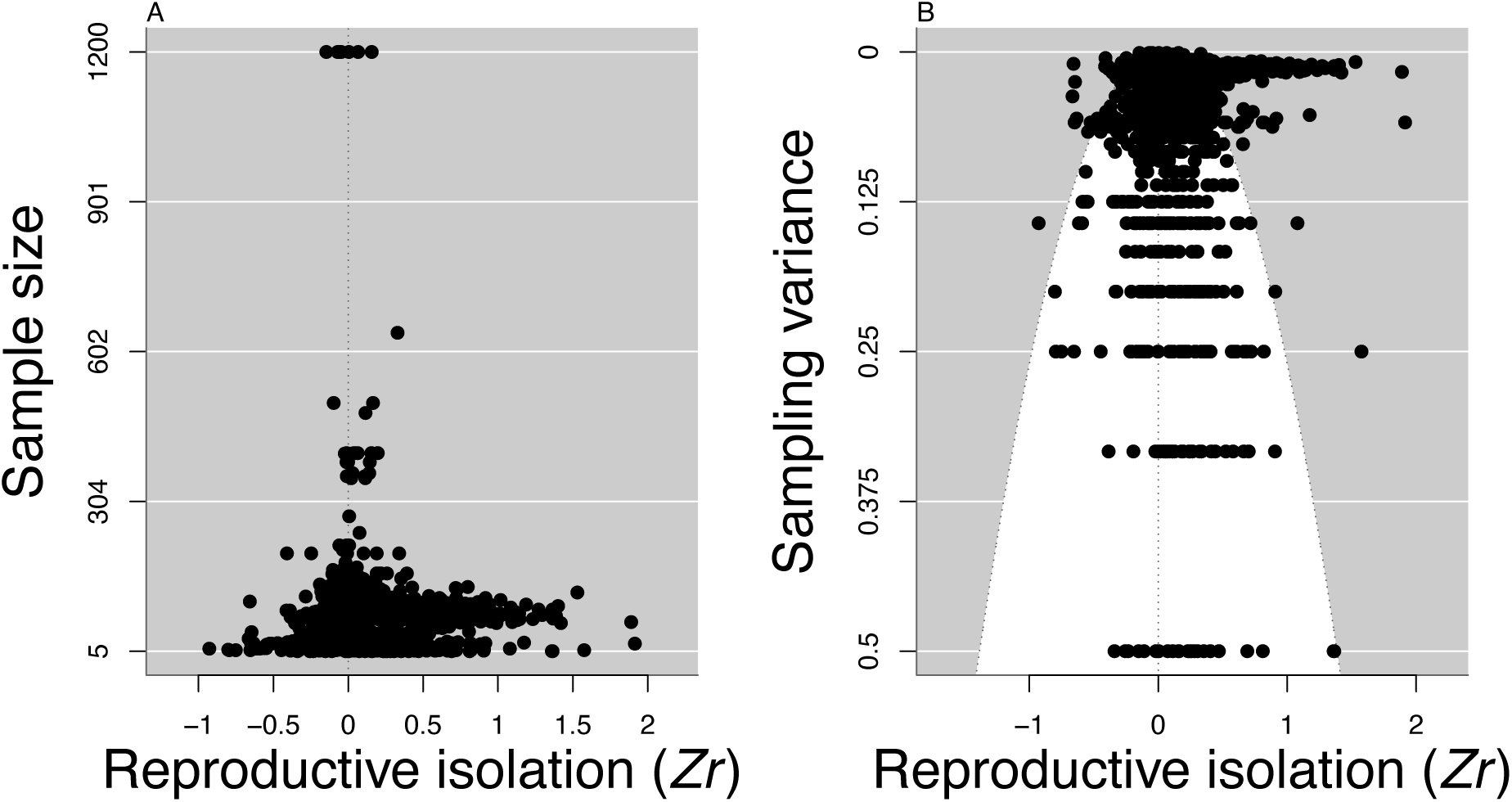
Meta-analytic funnel plots. The relationship between reproductive isolation and sample size (A) and reproductive isolation and sampling variance (B). Both plots show significant bias with skew to the right.

In our dataset, studies with smaller sample sizes reported larger effect sizes, indicating funnel asymmetry (Figure S5). Consistently, the slope of the relationship between our reproductive isolation effect sizes and their sampling variances was significant (Egger’s test: slope = 0.511, pMCMC = 0.03) in a multilevel meta-regression model accounting for repeated measures on different species and from different research groups, and non-independence due to phylogeny (Table S4). A trim and fill analysis indicated that 459 studies are needed to produce a symmetric funnel plot and the fail-safe number ranged from 1 723 to > one million, depending on the method of calculation. Note that the trim and fill analysis and the fail-safe numbers do not account for dependencies in the data. Finally, there was no evidence of a time-lag bias, which occurs when stronger effects are published more rapidly than weaker ones, in a multilevel meta-regression model accounting for dependencies in the data (slope = −0.001, pMCMC = 0.82). Taken together, these findings are indicative of small-study effects, which can result from publication bias (*118*).

There was considerable heterogeneity in our data set. In a random effects model not accounting for dependencies, I^2^ = 78%. In a multilevel meta-regression model accounting for dependencies, the I^2^ values were: repeated measures on different species = 2%; research group = 11%; phylogeny = 1%; residual = 61% (Table S4).

**Table S4.**
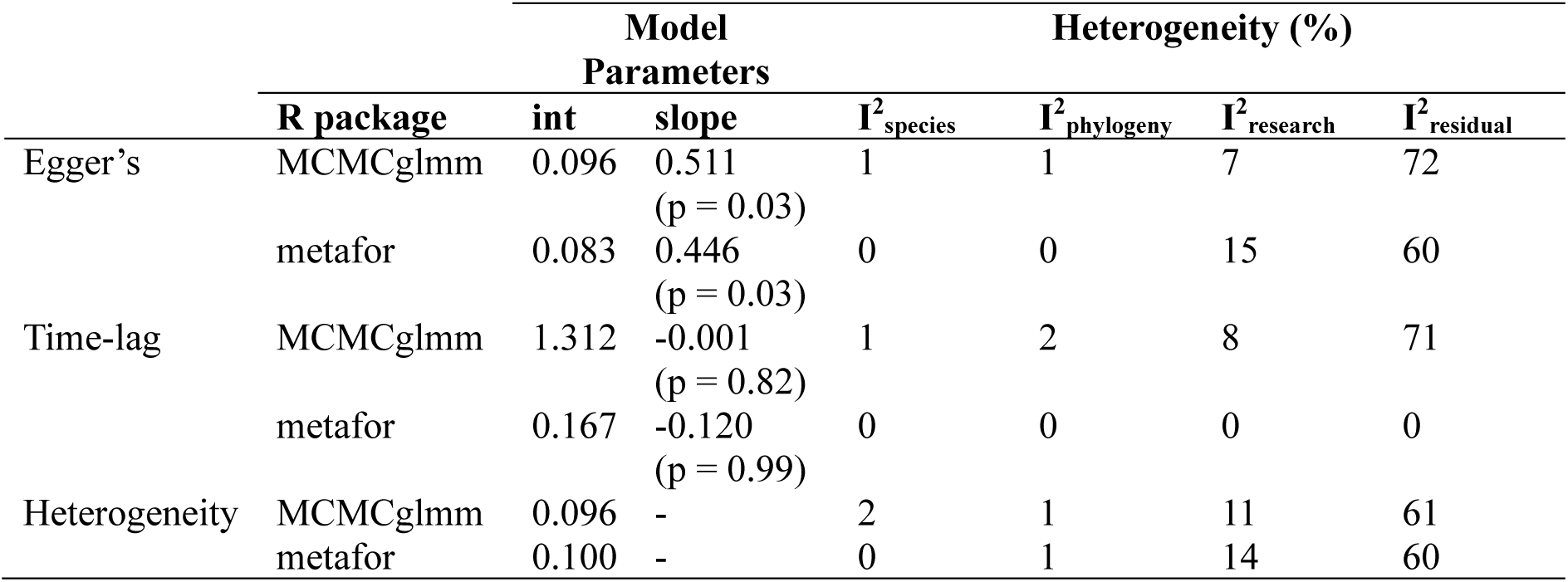
Results from publication bias and heterogeneity analyses performed using two statistical packages on the dataset without including any covariates.

#### 7.3 Main analysis

The main analyses were performed using both “MCMCglmm” (*119*) and “metafor” (*120*) in R (*111*). Both results are qualitatively and quantitatively similar, so we only presented the results from “MCMCglmm”. We used Inverse-Wishart priors for the random effects, and the default priors for fixed effects, with 1020000 iterations, 20000 of which were burnin, and a thinning interval of 1000.

We first fit a grand model that included: the cross type (either within environments or between environments); the number of generations at which reproductive isolation was estimated (ln-transformed); the barrier (pre- or post-mating); common garden (did the populations pass through a common garden, “Yes” or “No”); whether the data was at the population level (“Yes”, or estimates combined multiple populations, “No”); whether the data was backtransformed (“Yes” or “No”); and the founding population size of the populations used in the experiment (ln-transformed). To test for the interaction between time and cross-type, we fitted an interaction between them. As there was no support for the interaction, we removed it to report all other effects.

We included species, the species phylogeny, and the research group in which the work was completed as random terms. We included research group instead of paper ID as four studies measured reproductive isolation on two experimental evolution experiments (see Table S3 for more information). As we used multiple estimates of reproductive isolation from the same populations within experimental evolution studies, we used a variance (co)variance matrix of sampling variances that accounted for covariance between populations as the sampling variances for the model.

The analysis using “metafor” matched the results using “MCMCglmm” presented in the manuscript, but we display the “metafor” results in Table S5.

**Table S5.** Results of the main model from “metafor”. Table of coefficients for the main model, excluding the interaction between the number of generations and the between/within treatment variable.

|  | estimate | se | z-value | p-value | 95% CIs |
| --- | --- | --- | --- | --- | --- |
| Intercept | 0.116 | 0.105 | 1.113 | 0.266 | [-0.089, 0.321] |
| Between treatments | 0.067 | 0.016 | 4.167 | <0.001 | [0.035, 0.098] |
| Pre-mating barrier | 0.103 | 0.042 | 2.433 | 0.015 | [0.020, 0.186] |
| Common garden present | -0.087 | 0.041 | -2.114 | 0.035 | [-0.167, -0.006] |
| Number of generations (ln-transformed) | -0.023 | 0.016 | -1.450 | 0.147 | [-0.054, 0.008] |
| Population-level data | 0.167 | 0.053 | 0.323 | 0.747 | [-0.086, 0.120] |
| Backtransformed | -0.063 | 0.051 | -1.247 | 0.212 | [-0.163, 0.036] |
| Founding population size (ln-transformed) | 0.002 | 0.008 | 0.020 | 0.840 | [-0.013, 0.017] |

#### 7.4 Two-stage and random slope analysis

We found no effect of the number of generations a population evolved on reproductive isolation with the complete dataset. To more explicitly test this, we analysed only the studies that estimated reproductive isolation between the same populations more than twice. This only included five research groups (from seven papers, Figure S6), a drastically reduced dataset that prevents a detailed phylogenetically informed multilevel statistical approach. We ran a two-stage model in “metafor” (*120*), which combined parameters estimated from linear models fit for each research group separately in a mini-meta-analysis. Our purpose of this analysis was to focus on the evolution of reproductive isolation over time (generations) and so we did not investigate differences in slope for estimates of reproductive isolation between and within environments. We found support that the grand intercept was greater than 0 (estimate = 0.77 ± 0.38 se, 95% CIs = [0.02, 1.52], z = 2.02, p = 0.04), indicating that reproductive isolation did evolve in this subset of the data. However, we did not find evidence that the grand slope estimate was positive, or indeed different from 0 (slope estimate = −0.17 ± 0.12 se, 95% CIs = [−0.41, 0.06], z = −1.46, p = 0.15).

**Figure S6.**
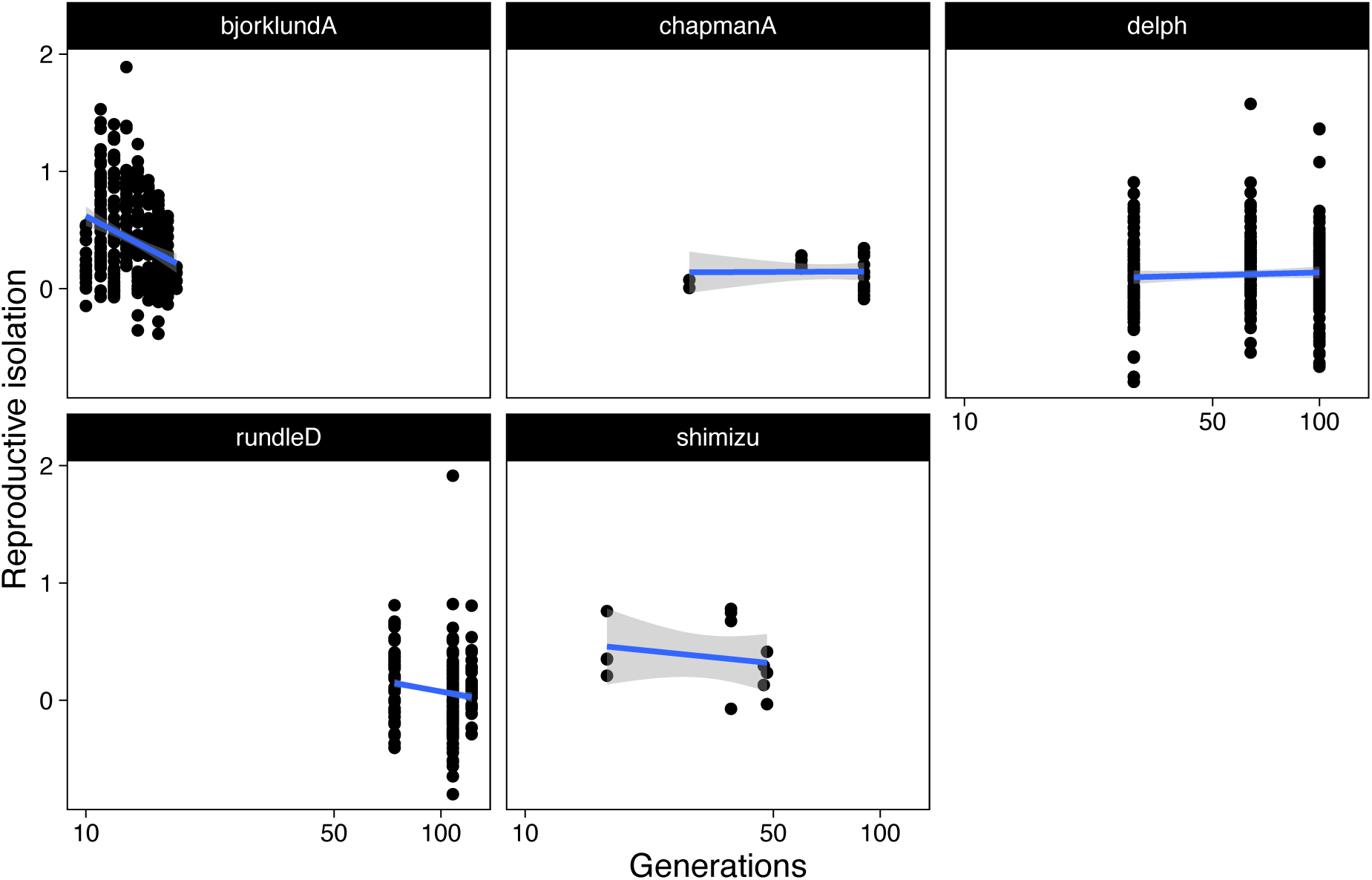
Studies that have estimated reproductive isolation in the same populations in more than one generation. Lines of best fit are shown to illustrate the overall patterns of reproductive isolation over time for each study as they are estimated without any reference to the covariance between populations used more than once to produce reproductive isolation estimates, or without reference to precision of the data. The papers that contributed data to each panel are: bjorklundA (107, 108); chapmanA (96, 102); delph (39); rundleD (68); and shimizu (100).

With the same dataset of five research groups, we ran a random slope model in “MCMCglmm” (*119*), which fits a random slope and intercept for each research group. In contrast to the two-stage model in “metafor” above, we found no evidence for an intercept that differs from 0 (estimate = 0.91, 95% Cis = [−0.24,2.22], pMCMC = 0.13). As above, this analysis also found no evidence that the number of generations changed the estimate of reproductive isolation (slope estimate = −0.22, 95% CIs = [−0.99,0.62], pMCMC = 0.47).

#### 7.5 Sensitivity analysis

In this section, we will outline a range of analyses to assess the robustness of the dataset. In many case, small decisions about which effect sizes to include, or which set of errors to include, may influence the overall results. This section is designed to explore some of these small decisions and their impact on our main results.

##### 7.5.1 Bayesian sampling variance

For Fisher’s z-transformed data of correlation coefficients, as our estimates of reproductive isolation effectively are being bounded between −1 and +1, 1 / n–3 is used as an estimate for the sampling variance for each effect size, where n is the sample size. We did, however, estimate reproductive isolation using Bayesian models in the R package “brms” to propagate errors from the data into the effect size. This approach also takes the sample size of the data into account, as it uses the actual data to estimate the standard deviation of the estimate, which we square for the sampling variance.

The use of the Bayesian sampling variance does not change the main results. We find: that reproductive isolation estimates between environments is greater than reproductive isolation within environments (difference estimate = 0.07, 95% CIs = [0.03, 0.10], pMCMC < 0.001); that pre-mating reproductive isolation estimates are greater than post-mating estimates of reproductive isolation (difference estimate = 0.14, 95% CIs = [0.07, 0.22], pMCMC < 0.001); that a common garden generation decreases the estimates of reproductive isolation (difference estimate = −0.11, 95% CIs = [−0.19, −0.02], pMCMC = 0.014); and that the number of generations (slope estimate = −0.004, 95% CIs = [−0.04, 0.03], pMCMC = 0.84), the founding population size (slope estimate = 0.005, 95% Cis = [−0.01, 0.02], pMCMC = 0.53), and whether the data were “backtransformed” (difference estimate = −0.08, 95% CIs = [−0.18, 0.01], pMCMC = 0.11) or from population-level crosses (difference estimate = 0.04, 95% CIs = [−0.02 ,0.11], pMCMC = 0.19) did not influence the evolution of reproductive isolation.

##### 7.5.2 Mean-centric estimates of reproductive isolation

Reproductive isolation was calculated a second way: using the average metric (mate choice proportion, hybrid fitness etc.) for each cross type and plugging it into

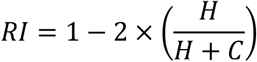

This would yield an estimate of reproductive isolation that includes no information about sample size or error.

The use of mean-centric reproductive isolation estimates also does not alter our main results, as coefficients are in the same direction and magnitude as the main analyses, even if the credible intervals just include zero in some cases. We find: that reproductive isolation estimates between environments is greater than reproductive isolation within environments (difference estimate = 0.06, 95% CIs = [0.02, 0.09], pMCMC < 0.001); that pre-mating reproductive isolation estimates are greater than post-mating estimates of reproductive isolation (difference estimate = 0.16, 95% CIs = [0.09, 0.23], pMCMC < 0.001); and that “backtransformed” data were more likely to yield estimates of reproductive isolation closer to 0 (difference estimate = −0.11, 95% CIs = [−0.19, −0.01], pMCMC = 0.03). The presence of a common garden generation did not influence estimates of reproductive isolation (difference estimate = −0.07, 95% CIs = [−0.15, 0.01], pMCMC = 0.11), despite the large parameter estimate that is similar to the main results. Similar to the main results, the number of generations (est = −0.01, 95% CIs = [−0.05, 0.02], pMCMC = 0.434), whether the data was from population-level estimates (difference estimate = −0.01, 95% CIs = [−0.07, 0.05], pMCMC = 0.79), and the founding population size (slope estimate = 0.002, 95% CIs = [−0.02, 0.02], pMCMC = 0.784) did not influence reproductive isolation.

##### 7.5.3 Exclusion of within-environment habitat isolation

Our metric of reproductive isolation varies between −1 and +1 except for within-environment estimates of habitat isolation. In this case, the metric of isolation is on which host plant females choose to oviposit. If two populations have been evolving on the same host plant and both choose to oviposit on that same host plant, reproductive isolation will equal 0. Estimates of reproductive isolation cannot go below 0. We therefore excluded these data in the analysis below and reanalysed the dataset. There are n = 78 estimates of within-environment habitat isolation that are excluded from this analysis.

Our results still stand for this analysis: reproductive isolation estimates between environments is greater than reproductive isolation estimates within environments (difference estimate = 0.08, 95% CIs = [0.05, 0.12], pMCMC < 0.001); pre-mating reproductive isolation estimates are greater than post-mating reproductive isolation estimates (difference estimate = 0.14, 95% CIs = [0.06, 0.21], pMCMC < 0.001); the number of generations does not influence reproductive isolation (slope estimate = −0.002, 95% CIs = [−0.03, 0.03], pMCMC = 0.908); the founding population size does not influence reproductive isolation (slope estimate = 0.006, 95% CIs = [−0.01, 0.02], pMCMC = 0.50); and whether the data are “backtransformed” does not influence reproductive isolation (difference estimate = −0.08, 95% CIs = [−0.19, 0.009], pMCMC = 0.096). There is less support for the result that a common garden decreases estimates of RI (difference estimate = −0.09, 95% CIs = [−0.17, 0.004], pMCMC < 0.001), though the parameter estimate is equivalent to the main analysis.

